# Intranasal dantrolene nanoparticles inhibit lipopolysaccharide-induced helplessness and anxiety behavior in mice

**DOI:** 10.1101/2024.09.06.611461

**Authors:** Jia Liu, Yan Lu, Piplu Bhuiyan, Jacob Gruttner, Lauren St. Louis, Yutong Yi, Ge Liang, Huafeng Wei

## Abstract

This study investigates the therapeutic effectiveness of intranasal dantrolene nanoparticles pretreatment to inhibit lipopolysaccharide (LPS)-induced pathological inflammation and synapse destruction and depressive and anxiety behavior in mice. B6SJLF1/J adult mice were pretreated with intranasal dantrolene nanoparticles (dantrolene: 5mg/kg), daily, Monday to Friday, 5 days per week, for 4 weeks. Then, mice were treated with intraperitoneal injection of LPS (5mg/kg) for one time. Behavioral tests for depression and anxiety were performed 24 hours after a one-time LPS injection. Biomarkers for pyroptosis-related inflammation cytokines (IL-1β and IL-18) in blood and brains were measured using enzyme-linked immunosorbent assay (ELISA) and immunoblotting, respectively. The changes of primary proteins activation inflammatory pyroptosis (NLRP3: NLR family pyrin domain containing 3, Caspase-1, N-GSDMD: N terminal protein gasdermin D) and synapse proteins (PSD-95 and synpatin-1) in brains were measured using immunoblotting. Intranasal dantrolene nanoparticles robustly inhibited LPS-induced depression and anxiety behavior. Intranasal dantrolene nanoparticles significantly inhibited LPS-induced pathological elevation of IL-1β and IL-18 in the blood and brain and inhibited LPS induced activation of pyroptosis. Intranasal dantrolene nanoparticles significantly ameliorated decrease of PSD-95 and synpatin-1 proteins in brains. Thus, intranasal dantrolene nanoparticles has demonstrated neuroprotection against inflammation mediated depression and anxiety behaviors and should be studied furthermore as a future effective drug treatment of major depression disorder or anxiety psychiatric disorder.

## Introduction

Three hundred million people worldwide suffer from major depressive disorder (MDD)^1^. MDD is a chronic and recurrent disease affecting about 20% of the population and the leading cause of suicide^2^. MDD imposes a tremendous psychological burden, as well as significant social repercussions and contributions to other disabilities^3^. The expected direct and indirect costs of MDD are up to $6 trillion in the USA alone^4^. An estimated 4% of the global population currently experience an anxiety disorder. In 2019, 301 million people in the world had an anxiety disorder, with a lifetime prevalence of approximately 34 percent^5^. Major depression disorder and anxiety disorders were approximately twice as prevalent among women^6,7^, with overall age-specific rates remaining relatively stable or increasing across the lifespan^8^. In 2021, an estimated 5.8 million (9.4%) children and adolescents were impacted by anxiety^9^. Depression and anxiety often co-exist and are the most common psychiatric diseases, which are chronic diseases with unclear mechanisms of pathology^10^.

Classical treatment of depression or anxiety psychiatric disorders includes atypical antipsychotics that modulate dopamine and serotonin neurotransmission^11,12^. Pharmacotherapies for anxiety disorders also include drugs that interact with GABA neurotransmission systems^13^. Unfortunately, antidepressants have significant limitations, including slow onset of action, high rates of nonresponse, and acute worsening of anxiety^14^. Benzodiazepines are not recommended for long-term use in some anxiety disorders, due to concerns about their potential for abuse, tolerance, and withdrawal, and they are ineffective in some anxiety spectrum disorders^15^. Treatment of MDD is prone to a high risk of resistance (up to 30% of patients are unresponsive to the first treatment) and relapse (up to 8%)^3,10^. Treatment-resistant anxiety with remission rates may be as low as 25–35%, and relapse rates post remission may be 30% after 10 years^16^. Thus, there is an urgent need to develop novel approaches to treat depression and anxiety, especially treatment-resistant depression, or anxiety, with minimal side effects or organ toxicity.

Increasing studies indicated that glutamate neurotransmission plays a critical role in mood function and its imbalance may cause psychiatric disorders^17^. Glutamate is linked to the development of anxiety disorders through its regulation of neuropeptides, fear extinction, and stress response. Glutamate is also important in synaptic and neural plasticity related to psychiatric disorders. Ketamine, an N-methyl-d-aspartate (NMDA) glutamate receptor antagonist, was approved in 1970 by the Food and Drug Administration (FDA) for use in children and adults as an anesthetic. Recent studies indicate that ketamine is effective in treating MDD, especially treatment-resistant depression (TRD)^18,19^. Ketamine is also effective in treating anxiety disorders, even those resistant to traditional treatments^9,20^. While traditional antidepressants can take weeks to months to have an effect, ketamine has rapid effects on mood and suicidality, with mood changes reported as early as the first 4 hours after treatment^21^. As an antagonist of NMDAR, a primary glutamate receptor on the plasma membrane that causes Ca^2+^ influx into the cytosol from extracellular space, ketamine has become a safe and broadly effective drug to treat depression or anxiety disorders, so is the intranasal esketamine^22^. This indicates that glutamate-mediated excitotoxicity and associated disruption of intracellular Ca^2+^ homeostasis play a key role in the pathology of depression and anxiety disorders.

Although molecular mechanisms are unclear, increasing evidence suggests that upstream disruption of intracellular Ca^2+^ homeostasis and associated downstream inflammation and synapse dysfunction play critical roles in MDD pathologies^23-25^, as well as anxiety disorder^26,27^. The overactivation of ryanodine receptors (RyRs) in MDD and associated excessive Ca^2+^ release from endoplasmic reticulum (ER) results in depletion of ER Ca^2+^ and pathological elevation of cytosol and mitochondrial Ca^2+^ concentrations, detrimental to synapse function and cell survival^25,28,29^. RyRs overactivation have also been shown to increase anxious behavior^28^. Upstream Ca^2+^ dysregulation results in mitochondria dysfunction^30^, mitochondrial and cellular oxidative stress^10,31^, activation of inflammasomes, and cell or neuron death by pyroptosis^32-35^. This results in the release of inflammatory cytokines (IL-1β and IL-18) and pathological inflammation-mediated cell death by pyroptosis^34,36^. Pathological cytokines, especially pyroptosis-related IL-18 play important roles in psychiatric disorders^37,38^. Gut dysbiosis and associated inflammation also contribute to pathology in depression and anxiety behaviors^39,40^. Thus, a drug that inhibits upstream Ca^2+^ dysregulation and then downstream pathological inflammation and programmed cell death by pyroptosis and synapse destruction is expected to treat MDD and anxiety disorder effectively.

Dantrolene, a RyRs antagonist, is a US Food and Drug Administration-approved drug for the treatment of malignant hyperthermia, muscle spasms, and neuroleptic syndrome, with tolerable side effects and occasional liver toxicity at high doses^41^. Dantrolene, with its ability to inhibit the common upstream critical Ca^2+^ dysregulation, is neuroprotective against many neurodegenerative diseases, including cerebral ischemia^42,43^, Huntington’s disease^44^, spinocerebellar ataxia^45^, amyotrophic lateral sclerosis^46^, and seizures^47,48^. Dantrolene robustly inhibited up to 78% of NMDA-induced elevation of cytosolic Ca^2+^ concentration in cerebral cortical neurons ^49^, suggesting it may have great potential to treat major depression disorder with a similar molecular mechanism of ketamine^50^, which needs to be studies urgently.

In this study, we investigated the therapeutic effectiveness and the potential mechanisms of intranasal dantrolene nanoparticles on depression and anxiety behavior associated with pathological inflammation in adult mice. We induced inflammation by intraperitoneal injection of LPS. Our results demonstrated that intranasal dantrolene nanoparticles for twelve consecutive weeks robustly inhibited LPS-induced depression and anxiety behaviors in adult mice. These beneficial effects of intranasal dantrolene nanoparticles treatment were associated with its robust and significant inhibition of LPS-induced pathological elevation of pyroptosis related inflammation cytokine of IL-1β and IL-18 in blood and brains and loss of synapse proteins (PSD-95 and synpatin-1) in brains. This study provides proof of concept that intranasal dantrolene nanoparticles may be an effective treatment for depression and anxiety psychiatric disorders, potentially by inhibiting pathological inflammation and synapse destruction in CNS.

## Materials and Methods

### Animals

All procedures were approved by the Institutional Animal Care and Use Committee (IACUC) at the University of Pennsylvania. All B6SJLF1/J adult mice were obtained from Jackson Lab. Mice were divided into different cages according to age and gender, with no more than five mice per cage. Both male and female mice were used in this study.

### Experimental treatment groups

As demonstrated in supplemental figure 1, adult male and female mice (5-10 months old) were randomly divided into different experimental groups. The mice were pretreated with either intranasal dantrolene nanoparticles (IN Dan, 5mg/kg) or vehicle (Veh), once a day, 5X/week (Monday through Friday). Control groups received no pretreatment. Fresh dantrolene nanoparticles were made every time before administration. Intranasal dantrolene nanoparticles stock solution was made at 5 mg/ml. The Ryanodex formulation vehicle was made fresh and contained all inactive ingredients in Ryanodex^51,52^. At the end of the 4-week pretreatment period, mice were then treated with one-time intraperitoneal injection of lipopolysaccharide (LPS, 5 mg/kg), once. Mice in the control group, without pretreatment, were treated with LPS (LPS, 5mg/kg) once as well. The sham control group received no pretreatment or treatment. At 24 hours following the one-time LPS treatment, behavioral tests for depression or anxiety behaviors were performed on all mice, with the following order of least to most stressful tests: OPT, EPMT, TST, and FST. Thereafter, all mice were euthanized, and the blood or brain were harvested for examination of inflammation and programmed cell death by pyroptosis and synapse destruction in blood and brains.

### Intranasal dantrolene nanoparticle or vehicle administration

Dantrolene (Sigma, St Louis, MO) was dissolved in the Ryanodex Formulation Vehicle (RFV: 125 mg mannitol, 25 mg polysorbate 80, 4mg povidone K12 in 5mL of sterile water and pH adjusted to 10.3), similar to our previous publications^52,53^. For intranasal administration, the final concentration of dantrolene was 5 mg/mL as we have previously described. Mice were held and fixed by the scruff of their necks with one hand. With the other hand, mice were given a total of 1μL/gram of body weight of dantrolene nanoparticles or RFV. Accordingly, a mouse weighing 20 g would be given 20μL of solution, which is equal to a dantrolene dose at dose at 5mg/kg. The solution was slowly delivered directly into the mouse’s nose. Care was taken to ensure that mice were minimally stressed, and that the solution remained in the nasal cavity and did not enter the stomach or lungs.

### Forced swimming test (FST)

Depression behavior was assessed using the FST, as described previously with some modification^54^. The mice from each group were settled in a clear glass tank of 25 cm height, 10 cm diameter, 15 cm water depth, and (23 ± 2) °C water temperature. The test’s total period was 6 minutes, 2 minutes for adaptation, and the total immobility time was recorded in the next 4 minutes. Immobility occurred when the mice discontinued floundering on the surface of the water, instead appearing as floating in an equilibrium state. The greater the immobility time, the more severe the depression behavior.

### Tail suspension test (TST)

Depression behavior was assessed using the tail suspension test described previously, with modifications^55^. We use specially manufactured tail suspension boxes, made of cartons with the dimensions (42 cm height X 18 cm width X 30 cm depth). To prevent animals from observing or interacting with each other, each mouse was suspended within its three-walled rectangular box. Each day, animals were acclimated to the testing room for at least 1 hour before the test. Each mouse was suspended in the middle of the box. The width and depth of the box were sufficiently sized to prevent the mouse from contacting the walls. In this setting, the approximate distance between the mouse’s nose and the apparatus floor was 20-25 cm. A plastic suspension bar (50 cm. height X 40 cm. depth), was used to suspend the tail of each mouse and positioned on the top of the box. At the bottom of each box, we placed a paper towel to collect feces or urine from the animals. A dark grey box, for albino animals, and a white colored box, for mice of other coat colors, were used. Before evaluation, the tail of the mouse was securely adhered to the suspension bar, which was able to withstand the mouse’s weight. A video camera was placed in position and started recording the TST session. The total duration of the test was 6 minutes. The paper towel was replaced after each trial. After all sessions were finished, Anymaze software (Stoelting, USA) was used to analyze the collected data.

During the behavioral analysis, the mobility time of each mouse was measured and subtracted from 360 seconds, producing the immobility time. The investigator was blinded to the animal groups. The greater immobility time, the more severe the depressive behavior.

### Open field test (OFT)

The OFT was performed as previously described to evaluate anxiety behavior as described^56^. Each mouse was placed individually into the OFT apparatus (44 × 44 × 44 cm3), facing the wall. The locomotor activity was recorded for six minutes with a camera above the apparatus under dim light. The anxiety behavior of each mouse was measured using the total distance traveled, center entries, immobility time, and time spent in the central zone. After each test, the apparatus was cleaned with 75% ethanol. Any-maze software (Stoelting USA) was used to analyze the collected data. The lesser the central zone distance and mean speed, the increased the anxiety behavior. The more immobile time, the more severe the anxiety behavior.

### Elevated plus maze test (EPMT)

Anxiety was also assessed using EPMT, as described previously, although with some modification^57^. Mice were placed in the center of an elevated plus-maze (arms are 33 cm × 5 cm, with 25 cm tall walls on the closed arms) under dim lighting and their behavior was videotaped for 5 minutes. We used Any-maze software (Stoelting USA) to analyze the collected data. The time spent in the closed and open arms, as well as the number of explorations of open arms, was measured and recorded. The more time spent in closed arms, the more severe the anxiety behavior.

### Euthanasia and tissue collection

Mice were euthanized after the completion of all behavioral tests. As described previously^52^, mice were deeply anesthetized with 2–4% isoflurane delivered through a nose cone, with the concentration was adjusted according to response to a toe pinch. The skin of each mouse was prepared, and an incision was made to open the chest and expose the heart. Blood was collected from the heart for the serum study using a syringe equipped with a 27G needle. The blood was centrifuged at 1400 rpm at 4°C for 30 min, and the supernatant was collected and frozen at –80°C. The mice were euthanized by transcardial perfusion and exsanguination was conducted with cold phosphate-buffered saline.

### Measurements of serum concentration of IL-1β and IL-18

We measured the serum IL-1β and IL-18 cytokines using an ELISA kit, following company instructions, and as we described previously for measurement of S100β, with some modification^58,59^. All reagents and samples (the supernatant of the blood) were thawed to room temperature (18 - 25 °C) before use. It is recommended that all standards and samples be run at least in duplicate. Add 100 μL of each standard and sample into appropriate wells. Cover wells and incubate for 2.5 hours at room temperature or overnight at 4 °C with gentle shaking. Discard the solution and wash it 4 times with 1X Wash Solution. Wash by filling each well with Wash Buffer (300 μL) using a multi-channel Pipette or auto washer. Complete removal of liquid at each step is essential to reliable performance. After the last wash, remove any remaining Wash Buffer by aspirating or decanting. Invert the plate and blot it against clean paper towels. Add 100 μL of 1x prepared Detection Antibody to each well. Cover wells and incubate for 1 hour at room temperature with gentle shaking. Discard the solution. Repeat the wash procedure as in step 3. Add 100 μL of prepared Streptavidin solution to each well. Cover wells and incubate for 45 minutes at room temperature with gentle shaking. Discard the solution. Repeat the wash as in step 3. Add 100 μL of TMB One-Step Substrate Reagent (Item H) to each well. Cover wells and incubate for 30 minutes at room temperature in the dark with gentle shaking. Add 50 μL of Stop Solution (Item I) to each well. Read absorbance at 450 nm immediately. Mouse IL-1 beta ELISA Kit and Mouse IL18 ELISA Kit come from the company SIGMA(RAB0275-1KT/RAB08100-1KT).

### Immunoblotting (Western Blot)

As we described previously^60^, total brain tissues were extracted by homogenization using cold RIPA buffer (#9806S, Cell Technology, USA) supplemented with protease inhibitor cocktails (P8340 Roche). The brain homogenates were rocked at 4°C for 90 minutes and then centrifuged at 14,000 rpm (Brushless Microcentrifuge, Denville 260D) for 20 minutes at 4°C to remove cell debris. After collecting the supernatant, the protein concentration was measured using a BCA protein assay kit (Pierce, Rockford, IL 61101 USA). Briefly, equal amounts of protein (20μg/lane) were loaded onto 4-20% gel electrophoresis of mini-protein TGX precast (Cat. #4561094, BIO-RAD) and transferred to polyvinylidene difluoride (PVDF) membranes (Immobilon-P, MERK Millipore, Ireland) using wet electrotransfer system (BIO-RAD, USA). Following the transfer membrane blocking with 5% BSA (Sigma-Aldrich) for 1 hour, then the PVDF membrane was incubated with primary antibodies overnight at 4 °C including NLRP3, cleaved caspase-1, GSDMDNT, IL-1β, IL-18, PSD95, Synapsin-1, GAPDH (Table 1). Then, the membranes were incubated with a secondary antibody including anti-Mouse IgG1 HRP-linked; Anti-rabbit IgG, HRP-linked that was conjugated to horseradish peroxidase and washed with Tris-buffered saline containing 0.2% Tween-20 (TBST). Following that, TBST was used for washing the membranes three time for 10 minutes. After incubation with secondary antibodies, protein bands were visualized using ECL Western Blotting Detection Reagents (Cytiva, amersham, UK) and quantified for band intensity using ImageJ software (National Institutes of health, Bethesda, MD, USA), by two persons blinded to treatments and averaged for each mouse.

**Table 1.**

|  | Cat. and Manufacture | Dilution |
| --- | --- | --- |
| Primary antibody |  |  |
| Anti- NLRP3 | 68102-1-Ig; Proteintech | 1:2000 |
| Anti-Human procaspase-1/ cleaved caspase-1 p20 | #2225S, Cell signaling Technology, USA | 1:1000 |
| Anti- Cleaved GSDMD | #36425S; Cell signaling Technology, USA | 1:1000 |
| Anti-IL-1β | 31202S, Cell signaling Technology, USA | 1:1000 |
| Anti-IL-18 | 10663-1-AP; Proteintech | 1:3000 |
| Anti-PSD95 | #2507; Cell signaling Technology, USA | 1:1000 |
| Anti-Synapsin-1 | #5297; Cell signaling Technology, USA | 1:1000 |
| GAPDH | MA5-15738, Thermo Fisher Scientific | 1:10,000 |
| Secondary antibody |  |  |
| Anti-Mouse IgG1; HRP -linked | # PA1-74421, Thermo Fisher Scientific | 1:10,000 |
| Anti-rabbit IgG, HRP-linked | #7074; Cell signaling Technology, USA | 1:1000 |

### Statistical Analysis

All data were represented as mean ± standard deviations (Means±SD). Statistical analyses were employed with GraphPad Prism (Version 9.3.1, CA, USA). Comparisons of more than two groups were conducted by one-way ANOVA with Tukey’s multiple comparison test. The N values in each group represent the number of mice. P<0.05 was considered statistically significant.

## Results

### Intranasal dantrolene nanoparticle pretreatment significantly inhibited LPS-induced helplessness and anxiety behaviors

The FST and TST tests are commonly used to evaluate helplessness or depression behaviors in mice and to test drug’s efficacy versus side effects^27,61,62^. In both FST (Fig. 1A) and TST (Fig. 1B), LPS induced significantly increased immobile times (helplessness or depression behaviors) by 89% (60.1 vs. 113.3) and 90% (98.1 vs. 186.6) respectively. Intranasal dantrolene nanoparticle pretreatment robustly inhibited the increased immobile time in FST by 41% (113.3 vs. 66.9) and in TST by 46% (186.6 vs. 101.5). In contrast, the intranasal dantrolene nanoparticles vehicle control did not demonstrate same inhibitory effects on helplessness behaviors. The elevated plus maze test (EPMT) and open field test (OPT) are commonly used to evaluate anxiety related behaviors in mice and test drug efficacy to treat anxiety behaviors^27,63,64^. In both EPMT (Fig. 1C) and OFT (Fig. 1D), LPS induced significantly increased time stay in closed arm in EPMT (anxiety behaviors) by 46% (162 vs. 237) and immobile time in OPF by 130% (82 vs. 189) respectively. Intranasal dantrolene nanoparticle pretreatment robustly inhibited the increased time stay in closed arm in EPMT by 29% (237 vs. 169) and immobile time in OPT by 40% (189 vs. 114). In contrast, the intranasal dantrolene nanoparticles vehicle control did not demonstrate same inhibitory effects on depression behaviors.

**Figure 1.**
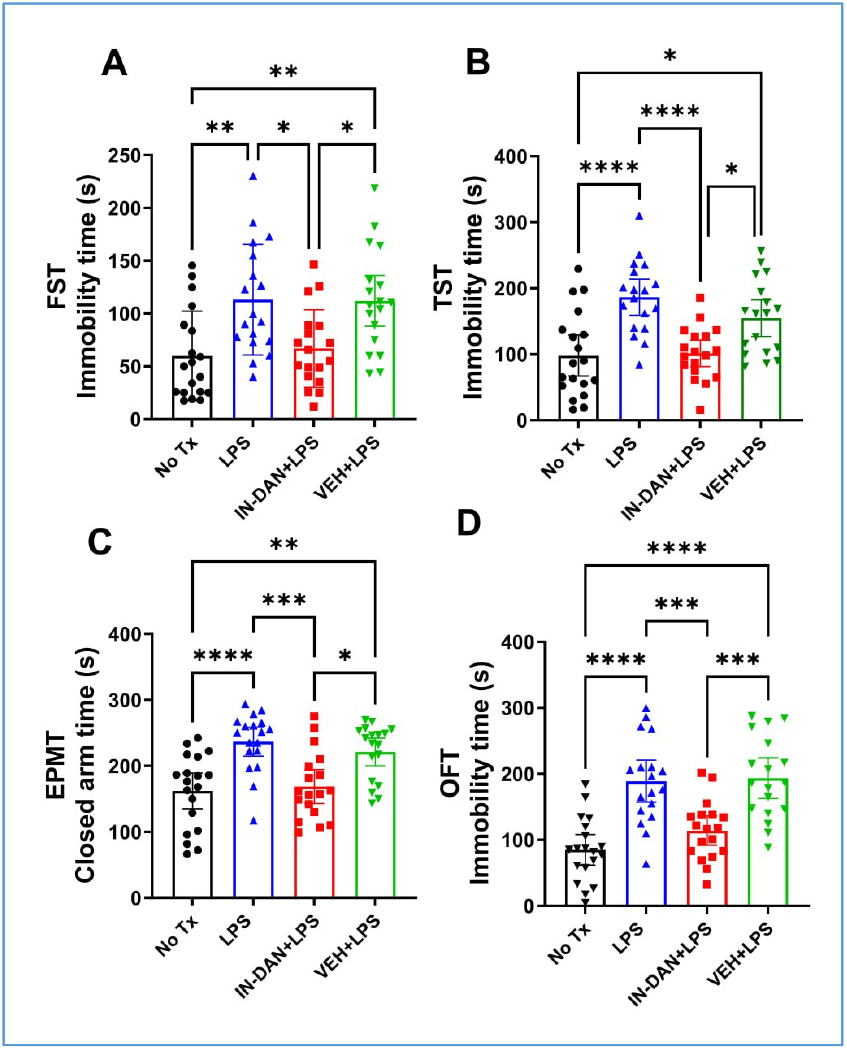
Intranasal dantrolene nanoparticles significantly reduced LPS-induced acute helplessness and anxiety-related behaviors. Adult B6SJLF1/J mice were treated with intranasal dantrolene nanoparticles (IN-DAN) or vehicle (VEH) for 28 days, followed by a single i.p. injection of lipopolysaccharide (LPS) (IN-DAN+LPS). Control mice either had no treatment (No Tx), only LPS (LPS), or VEH+LPS. The forced swim test (FST, A), tail suspension test (TST, B), elevated plus maze (EPMT, C) and open field test (OFT, D) were performed 24 hours after the LPS injection. Increased mobility time indicates helplessness or anxiety-related behaviors (A, B, D). Increased time in the closed arm indicates anxiety (C). N=18-19 mice, Mean±95%CI, One-Way ANOVA followed by Tukey post hoc test. *P<0.05, **P<0.01, ****P<0.0001

In the open field test, the less central zone distance mice traveled and less moving speed, the more severe the anxiety behavior (supplemental figure 2). Compared to control without treatment, one-time LPS treatment significantly decreased mean speed (anxiety behavior) but not traveled zone distance. Intranasal dantrolene nanoparticles trended to inhibit LPS induced decrease of travel zone distance and mean speed (supplemental figure 2).

### Intranasal dantrolene nanoparticle pretreatment significantly inhibited LPS-induced helplessness and anxiety behaviors primarily in female mice

We further analyze the effects of sex on neuroprotective effects of intranasal dantrolene nanoparticles on helplessness and anxiety behavior. As demonstrated in Fig. 2, intranasal dantrolene nanoparticles but not its vehicle control primarily inhibited helpless behavior determined by both FST (Fig. 2A) and TST (Fig. 2B) in female but not male mice. For anxiety behavior, although intranasal dantrolene nanoparticle but not vehicle control inhibited LPS induced anxiety behavior determined by EPMT (Fig. 2C) in male mice, the inhibitory effects of dantrolene on anxiety behaviors were most significant in female mice determined by both EPMT (Fig. 2C) and OFT (Fig. 2D).

**Figure 2.**
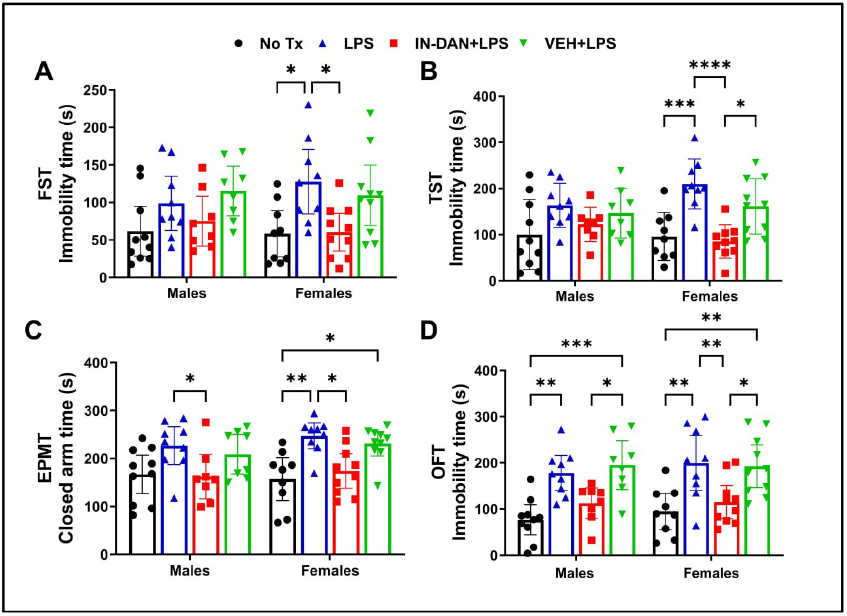
Intranasal dantrolene nanoparticles reduced LPS-induced acute helplessness and anxiety-related behaviors more in female than male mice. The sex of the adult B6SJLF1/J mice, pretreated with intranasal dantrolene nanoparticles (IN-DAN) or vehicle (VEH) for 28 days, followed by a single i.p. injection of LPS (IN-DAN+LPS), was examined. No treatment controls (No Tx), LPS only (LPS), or VEH+LPS. The forced swimming test (FST, A), tail suspension test (TST, B), elevated plus maze (EPMT, C), and open field test (OFT, D) were performed 24 hours after the LPS injection. N=8-10 mice, Mean±95%CI, Two-Way ANOVA followed by Tukey post hoc test. *P<0.05,**P<0.01, ****P<0.0001

### Intranasal dantrolene nanoparticles significantly inhibited LPS-induced pathological elevation of pyroptosis-related inflammation cytokines in blood and brains

Compared to control without treatment, IP injection of LPS (5mg/kg) for one time significantly increased blood concentrations of IL-1β by 147% (Fig. 3A, 136 vs. 336) and IL-18 by 67% (Fig. 3B, 233 vs. 390). Intranasal dantrolene nanoparticles but not its vehicle control significantly inhibited the increased IL-1β by 46% (336 vs. 183) and elevated IL-18 by 23% (Fig. 3B, 390 vs. 299). Furthermore, intranasal dantrolene nanoparticles significantly inhibited LPS-induced elevation of IL-1β and IL-18 proteins in both blood (Fig. 3 A, B) and in brains (Fig. 3 C, D, supplemental figure 3 and 4).

**Figure 3.**
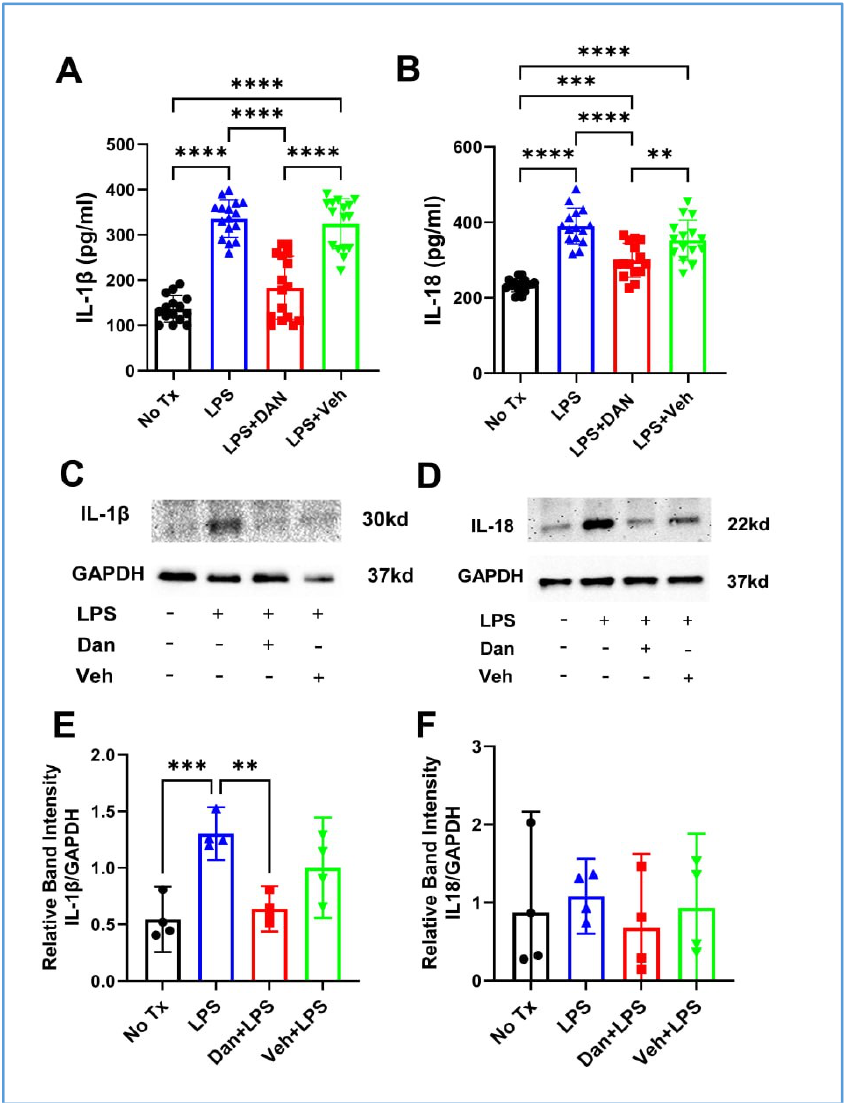
Intranasal Dantrolene Significantly Inhibited lipopoly-.saccharide-induced pathological elevation of IL-18 and IL-1β cytokines in the blood and brains. B6SJLF1/J adult mice were pretreated with intranasal dantrolene nanoparticles DAN, 5mg/kg) or vehicle (Veh) control daily, Monday to Friday, for continuous 4 weeks. Mice were then treated with one-time IP injection of lipopolysaccharide (LPS, 5 mg/kg). Mice in control (No Tx) group received no treatments. IL1β in blood (A) and brain (C, E) and IL18 in blood (B) and brains (D, F) were measured using ELISA assay kit (A, B) or immunoblotting (C-F)., N=18-20 mice for blood measurement (A, B) and N=4 brains for immunoblotting (C-F), Means±95% CI, One-Way ANOVA followed by Tukey post hoc test. **P<0.01, ****P<0.0001 respectively.

### Intranasal dantrolene nanoparticles inhibited LPS-induced programmed cell death by pyroptosis in mice brains

Compared to control mice without treatment, LPS treatment increased the proteins levels of pyroptosis pathway activation (NLRP3, caspase-1 and N terminal GSDMD) in mice brains. Intranasal dantrolene nanoparticles significantly inhibited increase of Caspase-1 (P20), a biomarker of programmed cell death by pyroptosis (Fig. 4 C, D, supplemental figure 6), and trended to inhibit LPS-induced elevation of pyroptosis pathway activation proteins (Figure 4, A and B, supplemental figure 5, NLRP-3 and Fig. 4 E and F, supplemental figure 7, N terminal GSDMD (GSDMD-NT))

**Figure 4.**
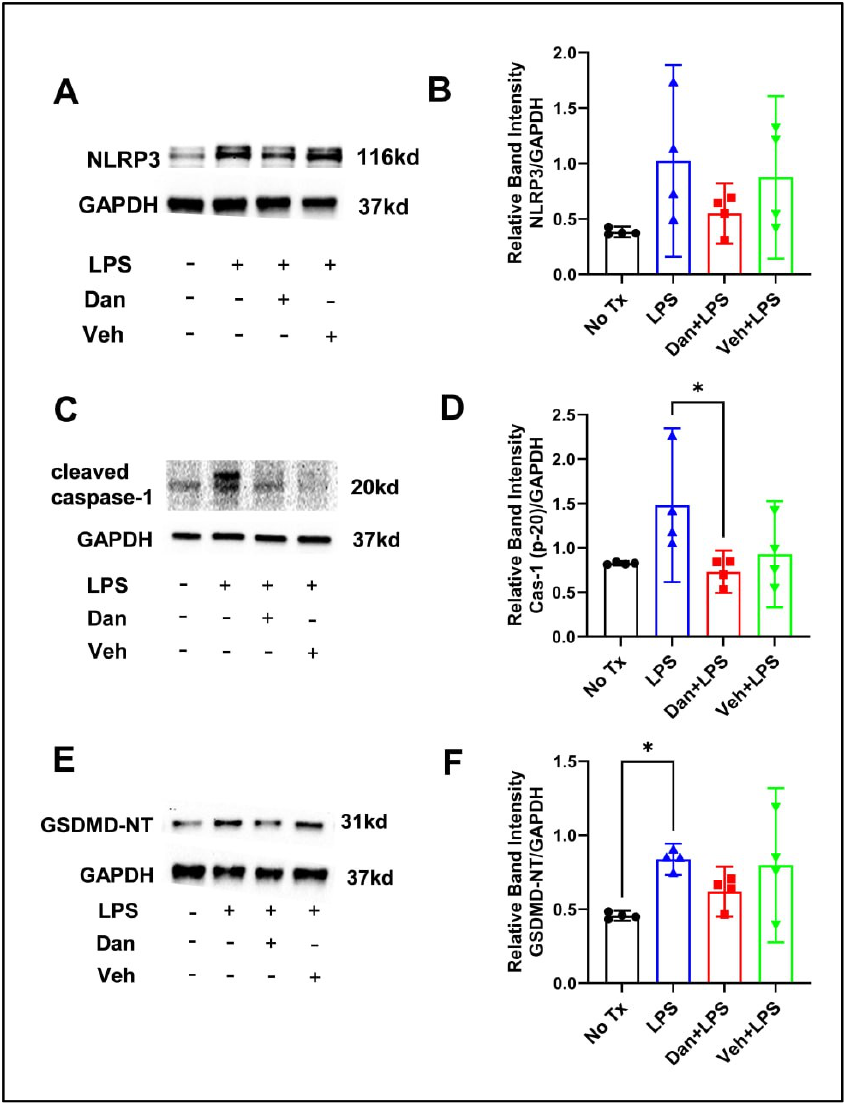
Effects of intranasal dantrolene nanoparticles on lipo-polysaccharide-induced programed cell death by pyroptosis in the brains. B6SJLF1/J adult mice (5-10 months old) were pretreated with intranasal dantrolene nanoparticles (Dan, 5mg/kg) or vehicle (Veh) daily, Monday to Friday, for continuous 4 weeks. Mice were then treated with one-time IP injection of lipopolysaccharide (LPS, 5 mg/kg). Mice in control (No Tx) group received no treatments. Representative western blot (A, C, E) and statistical analysis (B, D, F) determined the protein levels of critical regulatory proteins of pyroptosis pathway (A, B, NLRP3; C, D, Active caspase-1 (P-20), E, F, N terminal GSDMD (GSDMD-NT) in brains. N=4 different mice brain in each group. GAPDH as loading control. Means±95%CI, One-Way ANOVA followed by Tukey post hoc test. * P<0.05.

### Intranasal dantrolene nanoparticles significantly inhibited LPS-induced synapse protein loss in mice brains

Compared to control without treatment, LPS significantly decreased the synapse proteins of PSD-95 by 45% (1.322 vs. 0.726, Fig. 5, A, C, supplemental figure 8) and synapsin-1 by 37% (0.925 vs. 0.579, Fig. 5, B, D, supplemental figure 9) in mice brains. Intranasal dantrolene nanoparticles but not vehicle control pretreatment for 4 weeks significantly inhibited the pathologically reduction of synapse proteins of PSD-95 by 77% (0.726 vs. 1.286, Fig. 5 A, C, supplemental figure 8) and synpasin-1 by 76% (0.579 vs. 1.021, Fig. 5 B and D, supplemental figure 9).

**Figure 5.**
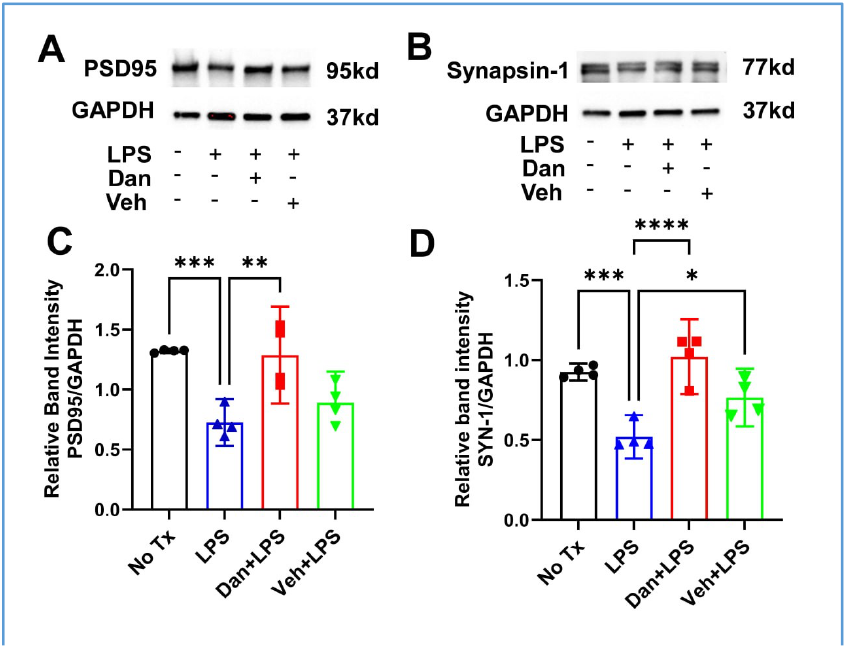
Intranasal dantrolene nanoparticles significantly inhibited lipopolysaccharide-induced synapse protein loss in brains. B6SJLF1/J adult mice were pretreated with intranasal dantrolene nanoparticles (Dan, 5mg/kg) or vehicle (Veh) daily, Monday to Friday, for continuous 4 weeks. Mice were then treated with one-time IP injection of lipopolysaccharide (LPS, 5 mg/kg). Mice in control (No Tx) group received no treatments. Western blot measured changes of synapse proteins of PSD-95 (**A, C**) and Synapsin-1 (**B, D**) in the brains. GAPDH as loading control. N=4 different mice in each group, Means±95%CI, One-Way ANOVA followed by Tukey post hoc test. *P<0.05, **P<0.01, ***P<0.001, P<0.0001 respectively.

## Discussion

There is an urgent need for new therapeutic interventions to prevent or treat MDD^3,65^ and anxiety5 psychiatric disorders, especially those drug-resistant and recurrent depression and anxiety^5,16,66^. This study has demonstrated that intranasal dantrolene nanoparticles are capable of ameliorating helplessness and anxiety behaviors in adult mice, associated with its ability to inhibit pathological elevation of pyroptosis related cytokines (IL-1β and IL-18) in blood, pyroptosis biomarker caspase-1 (P20) and synapse proteins loss in brains. Considering that upstream Ca^2+^ dysregulation plays an important role in the pathology of depression and anxiety, evidenced by the effectiveness of ketamine and esketamine in treating depression and anxiety resistant to traditional drug treatments^14,16^, a new class of drugs, such as dantrolene, could be developed with a mechanism to correct the disruption of upstream Ca^2+^ dysregulation and associated downstream pathological inflammation and pyroptosis in psychiatric disorders.

Increasing studies suggests that inflammation is an important pathology causing MDD and anxiety and could be a target for effective treatments^67-70^. Intracellular Ca^2+^ dysregulation has been proposed as the upstream cause of downstream mitochondria dysfunction, oxidative stress, and inflammation in MDD^25,71,72^ and anxiety^38,73^. Inhibition of upstream Ca^2+^ dysregulation through inhibiting over-activation of RyRs has been shown to suppress inflammation in Gulf War illness characterized by depression and dementia^28^. Although dantrolene has been shown to inhibit pathological inflammation with increased cytokines in diabetes^74,75^, sepsis^76,77^ and in COVID-19 infection^78,79^, its effects to inhibit Ca^2+^ dysregulation and associated inflammation for the treatment of depression and anxiety behaviors has not yet been investigated until now and studied herewith.

Several animal models to establish depression and anxiety behaviors have been suggested based on the proposed neurobiology mechanisms, including altered neurotransmission, HPA axis abnormalities involved in chronic stress, inflammation, reduced neuroplasticity, and network dysfunction^80-82^. Induction of inflammation in animals has been increasingly used to establish depression and anxiety animal models, especially using LPS to induce inflammation, depression, and anxiety behaviors to test drug treatment efficacy^37,71,83-86^. The common approach of LPS administration is intraperitoneal injection because of its easiness and feasibility. The dose and duration of IP injection of LPS to establish depression or anxiety behaviors has been varied but appears to be dose dependent. The high dose of LPS at 5 mg/kg, administered once, seems to be dependable in inducing inflammation and establishing depression behavior71. Our study confirmed the efficiency of a one-time administration of IP LPS (5 mg/kg) to induce inflammation and depression and anxiety behavior in adult mice consistently. Accordingly, the LPS-induced inflammation-mediated depression behaviors are adequate to evaluate the therapeutic effects of dantrolene to treat depression. Our results demonstrated robust inhibition of LPS-induced inflammation and associated helplessness and anxiety behaviors by intranasal dantrolene nanoparticles pretreatment in adult mice. In addition with the protective effects of dantrolene against memory loss in various AD animal models^52,87-89^, intranasal dantrolene nanoparticles also significantly inhibited helplessness and anxiety behaviors, and associated inflammation and pyroptosis, suggesting that pathological inflammation and pyroptosis play a critical role in both cognitive dysfunction and psychiatric disorders. Drugs that target inflammation pathology should be developed to treat both memory loss and psychiatric disorders in AD patients. Nevertheless, intranasal dantrolene nanoparticles significantly inhibited LPS-induced pathological elevation of pyroptosis related inflammation cytokines (IL-1β and IL-18) in blood, and ameliorated both depression and anxiety behavior in adult mice, suggesting that intranasal dantrolene nanoparticles could be developed as an effective drug to treat depression and/or anxiety in major depression disorder (MDD) patients potentially by its inhibition of inflammation and associated programmed cell death by pyroptosis.

MDD affect female twice more than male patients^90,91^. The proposed mechanisms for this sex difference include the effects of estrogen^92^, serotonin and tryptophan metablism93, different reactions to stresses^91^ etc. An important finding in this study is that intranasal dantrolene nanoparticles primarily be effective to ameliorate helplessness and anxiety behaviors primarily in female than in male mice. Our result is consistent with MDD prevalence, more in female than male mice. Also, the more significant inhibition of inflammation associated helplessness and anxiety behavior in female mice by intranasal dantrolene nanoparticles suggested that dantrolene may inhibit critical pathology and etiology contributing depression and anxiety behaviors. Although the exact mechanism unclears, we propose that dantrolene inhibited the upper stream critical Ca^2+^ dysregulation and associated inflammation and associated programmed neuron death by pyroptosis and synapse protein loss.

Intranasal administration of drugs, especially in a nanoparticle formulation, significantly promotes drugs that bypass the blood-brain barrier (BBB) and penetrate the CNS, with reduced peripheral toxicity^94-96^. Our recent research work indicated that intranasal dantrolene nanoparticles achieved higher therapeutic efficacy in the brain compared to oral and subcutaneous forms administration, with minimal or no side effects after chronic use^52,53,97^. Intranasal dantrolene nanoparticles significantly ameliorated memory loss in adult 5XFAD mice as a disease-modifying drug^52^. Increasing studies suggest ryanodine receptor overactivation and associated Ca^2+^ dysregulation is an upstream critical pathology leading to multiple downstream pathologies including mitochondria dysfunction^98^, oxidative stresses^99^, pathological inflammation^100^, and neuron damage by pyroptosis^101^. Eventually, these pathologies result in depression and/or anxiety psychiatric disorders^34,70,102^. This study demonstrated that inhibition of RyRs overactivation by dantrolene inhibited pathological inflammation and synapse proteins destruction and ameliorates depression and anxiety behavior robustly. This study further strengthens the indication that upstream RyRs overactivation and Ca^2+^ dysregulation and associated downstream pathological inflammation and synapse destruction play important roles in depression and anxiety psychiatric disorders. Like esketamine^22,103^, the intranasal dantrolene nanoparticles could become an effective disease-modifying drug treatment for both depression and anxiety psychiatric disorders and need to be investigated in future clinical studies.

Since depression and anxiety psychiatric disorders are chronic diseases, they require long-term drug treatments. Accordingly, proposed drugs must limit side effects or organ toxicity with chronic treatment. The major advantage of using intranasal dantrolene nanoparticles, in comparison to commonly used oral or intravenous approaches, is that it significantly increases the brain/blood concentration ratio of dantrolene^52,53^, especially in aged mice^97^. This promotes its CNS therapeutic effects, while minimizing its side effects or organ toxicity^52^. Our previous study demonstrated no side effects on nose structure and smell function^52,104^, liver structure^52^ and function, or muscle function^51,52,87^ after up to 10 months of chronic treatment in adult 5XFAD mice^52^, suggesting that intranasal dantrolene nanoparticles are safe to be used chronically in animals. Chronic use of dantrolene in patients’ needs to be investigated further.

This study has the following limitations: 1). The mice age is from 5 to 10 months old. Although they are considered adult mice, future studies need to focus on different age group, especially in aged mice, considering the high prevalence of psychiatric disorder s in aged population. 2). We were unable to measure the changes of cytosol versus mitochondria Ca^2+^ levels in the brain tissue due to technological challenges. However, dantrolene has been shown to inhibit LPS or AD gene mutation-induced overactivation of RyRs and associated Ca^2+^ dysregulation in different cell culture models^105,106^. 3). We did not measure the contents of reactive oxygen species (ROS) concentrations in brains which is usually the upstream pathology to activate NLRP3 inflammasome^107^. 4). We did not measure other biomarkers demonstrating effects of intranasal dantrolene nanoparticles on programed cell death by pyroptosis in brains. 6). We did not study the dose-response of dantrolene to treat depression and anxiety behaviors. Physiological Ca^2+^ release from ER is important for many physiological functions in cells, which means over-inhibition of calcium release from the ER may be harmful to cells, as we described previously. The adequate dose of intranasal dantrolene nanoparticles to treat psychiatric disorders must be investigated thoroughly before its clinical use in patients.

In conclusion, this study demonstrated that intranasal dantrolene nanoparticles significantly ameliorated inflammation mediated depression and anxiety behaviors in adult mice. This neuroprotective effect against depression and anxiety behaviors was associated with its robust inhibition of pathological elevation of pyroptosis related inflammation cytokines in blood and the synapse destruction in brains. This study may inspire future studies to repurpose intranasal dantrolene nanoparticles in treating depression and anxiety psychiatric disorders.

## Acknowledgements

This work was supported by grants to HW from the National Institute on Aging (R01AG061447) and NIA R01 Supplemental (3R01AG061447-03S1). The research was performed in the lab of Dr. Huafeng Wei and should be attributed to the Department of Anesthesiology, University of Pennsylvania. We appreciate the technical support from Rebecca Chae from Rowan University, New Jersey, U.S.A and Kyulee Kim from UPENN, Pennsylvania, U.S.A.

## Author contributions

H.W. conceived and designed the study. J.L., Y.L, P.B. J.G, L.S.L, Y.Y., G.L, H.W. conducted experiments, acquired and the data, J.L., Y.L, P.B. J.G, L.S.L, Y.Y., G.L, H.W. analyzed data and contributed to the manuscript preparation H.W. wrote the manuscript. All the authors reviewed and approved the final manuscript.

---

**Supplemental figure 1.**
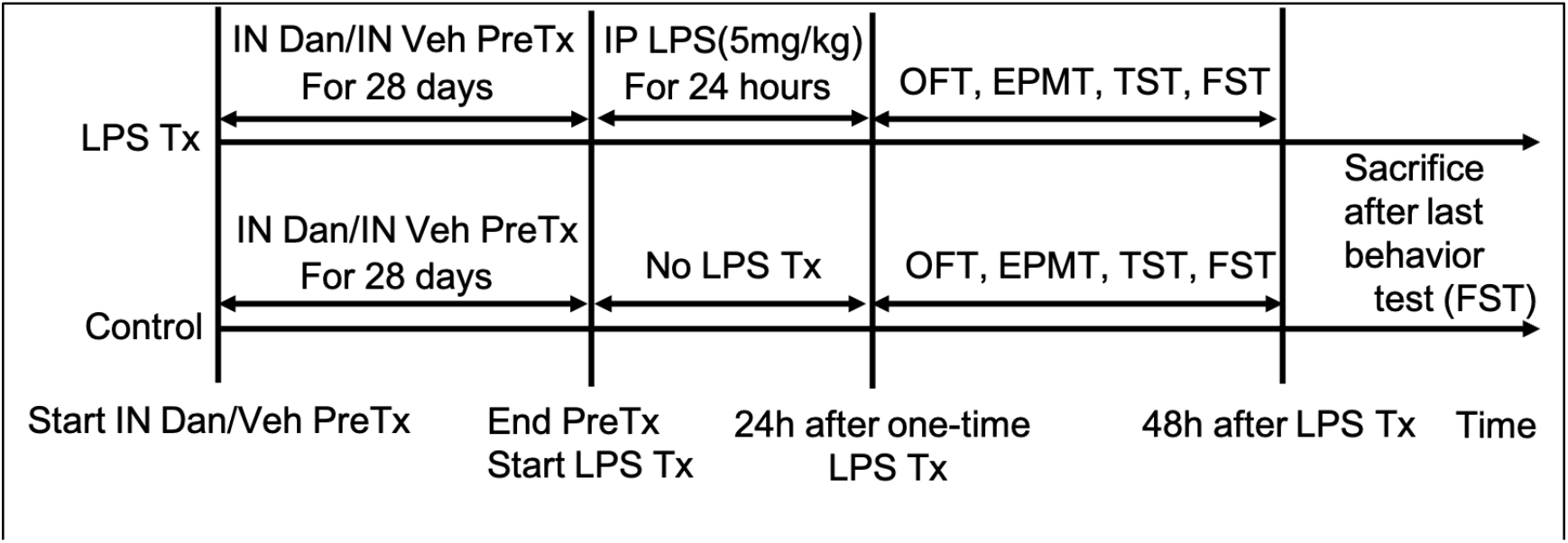
Experimental Design. Adult B6SJLF1/J mice were pretreated with intranasal dantrolene nanoparticles in Ryanodex formulation (DNRF, 5mg/kg) or vehicle (Veh) for 4 weeks. Control group received no treatments. Mice were then treated with intraperitoneal (IP) injection of lipopolysaccharide (LPS, 5mg/kg) for one time. Depression and anxiety behavior tests were performed at 24 hours after one-time IP LPS injection. PreTX: Pretreatment, Tx: Treatment, IN: Intranasal, Dan: Dantrolene, OFT: Open Field Test, EPMT: Elevated Plus Maze Test, TST: Tail Suspension Test, FST: Forced Swimming Test

**Supplemental figure 2.**
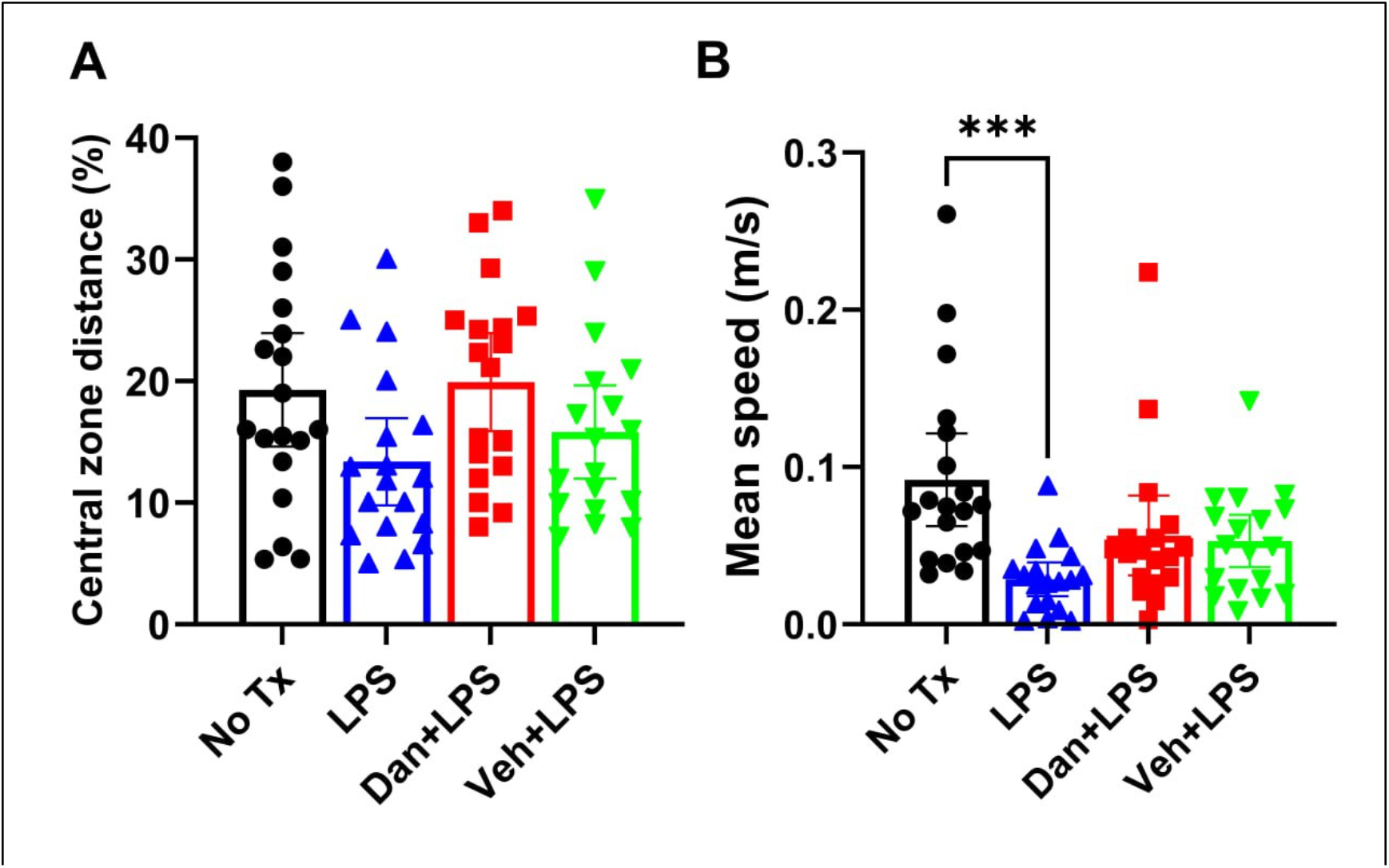
Effects of intranasal dantrolene nanoparticles on lipopolysaccharide-induced anxiety behavior in adult mice. B6SJLF1/J adult mice (5-10 months old) were pretreated with intranasal dantrolene nanoparticles (Dan, 5mg/kg) or vehicle (Veh) control daily, Monday to Friday, for continuous 4 weeks. Mice were then treated with one-time IP injection of lipopolysaccharide (LPS, 5 mg/kg). Mice in control (No Tx) group received no treatments. Open field test (OFT) was performed at 24 hours after one-time IP LPS injection. The less central zone distance or less mean speed, the more anxiety behavior for OFT. N=18-20 mice. Means±95%CI, One-Way ANOVA followed by Tukey post hoc test. ***P<0.001.

